# Genome-wide CRISPR guide RNA design and specificity analysis with GuideScan2

**DOI:** 10.1101/2022.05.02.490368

**Authors:** Henri Schmidt, Minsi Zhang, Haralambos Mourelatos, Francisco J. Sánchez-Rivera, Scott W. Lowe, Andrea Ventura, Christina S. Leslie, Yuri Pritykin

## Abstract

We present GuideScan2 for memory-efficient, parallelizable construction of high-specificity CRISPR guide RNA (gRNA) databases and user-friendly gRNA/library design in custom genomes. GuideScan2 analysis identified widespread confounding effects of low-specificity gRNAs in published CRISPR knockout, interference and activation screens and enabled construction of a ready-to-use gRNA library that reduced off-target effects in a novel gene essentiality screen. GuideScan2 also enabled the design and experimental validation of allele-specific gRNAs in a hybrid mouse genome.

## Main Text

CRISPR-based technologies have transformed life sciences and shown promise in the development of therapeutics [1], and genome-wide CRISPR screens are routinely used for the unbiased identification of regulators of a diverse range of cellular phenotypes. However, the design of efficient and specific guide RNAs (gRNAs) for CRISPR-based genomic perturbations presents computational challenges. Unwanted gRNA off-targets can cause inefficient targeting as well as genotoxicity, and incomplete information about off-targets can result in misinterpretation of experimental results [2]. We previously developed GuideScan [3] for scalable gRNA design, and we and others have demonstrated that GuideScan is more accurate than other tools in enumerating potential off-targets and estimating gRNA specificity [3, 2]. A key observation was that short-read aligners used by other gRNA design tools, while highly efficient for typical read count quantification tasks, do not exhaustively count suboptimal alignments or even multiple perfect alignments (without mismatches) [3]. GuideScan overcame this limitation by using a custom retrieval tree (trie) data structure to preprocess and analyze the targetable genome. The trie is constructed and stored in memory and then used to enumerate all off-targets of all potential gRNAs, thereby assessing their specificity. The database is then constructed from all uniquely targeting gRNAs. Despite addressing the main limitation of previous gRNA design tools, GuideScan requires that the targetable space is pre-specified: for example, for CRISPR-Cas9, only 20-nucleotide long sequences (20-mers) followed by the protospacer-adjacent motif (PAM) sequence NGG are considered as primary targets. While the resulting gRNA database is compactly stored on disk and allows for efficient access by genomic coordinates, the intermediate GuideScan trie data structure requires a large amount of memory for mammalian genomes (e.g. *>*190Gb for hg38), limiting the ability to manipulate the data structure or parallelize the gRNA database construction. For example, performing specificity analysis on gRNAs that are not in the GuideScan database would require loading this large trie data structure into memory.

Here we present GuideScan2, a new gRNA design and analysis software that overcomes the limitations of GuideScan and significantly expands its functionality. GuideScan2 implements preprocessing of an entire input genome into a lightweight index based on a compressed Burrows-Wheeler Transform (**Fig. 1, Methods**). Therefore, GuideScan2 is built upon the same efficient data structure as short-read aligners but is engineered to exactly account for off-targets. This approach does not require prespecifying the targeting rules, and the same index can be used for analysis and database construction for different gRNA lengths, PAM sequences, and off-target definitions. The GuideScan2 index is memory-efficient (e.g. 3.4Gb for hg38) and fast to construct (*≈* 30 min on a standard laptop), enabling efficient online access and scalable search by gRNA sequence. At the same time, the resulting gRNA database is as accurate as GuideScan in enumerating all potential gRNA off-targets and estimating off-target specificity. GuideScan2 functionality for processing the genome, for index and database construction and manipulation, as well as for gRNA design and analysis, is available through an open source command-line software package (https://github.com/schmidt73/guidescan-cli). We also provide a user-friendly web interface for gRNA design and analysis at https://guidescan.com.

**Figure 1:**
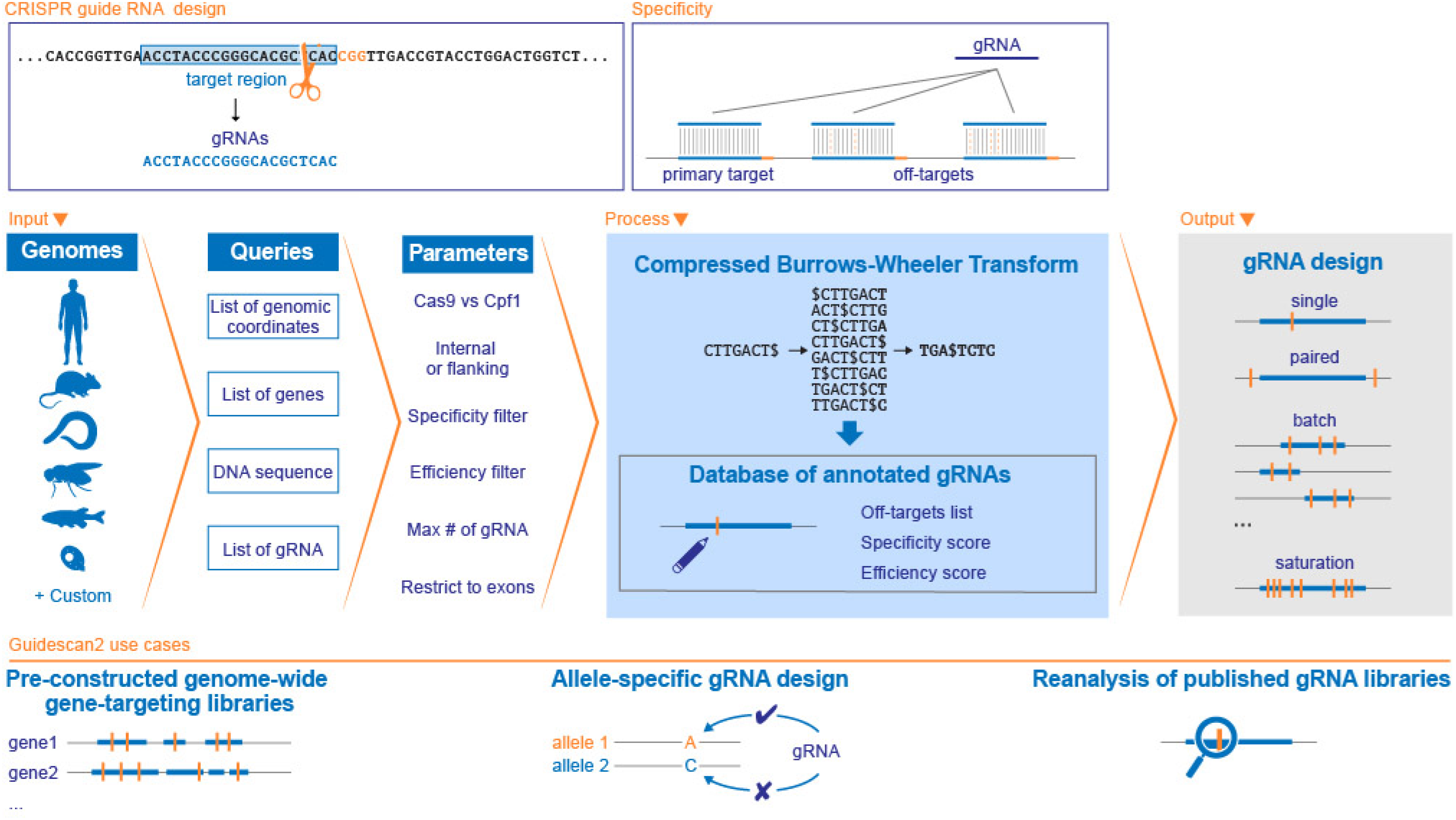
Overview of GuideScan2. GuideScan2 enables the design of gRNAs or gRNA libraries while exactly accounting for off-target effects. GuideScan CRIPSR-Cas9 and -Cas12a (Cpf1) libraries are precomputed for standard genomes; databases for custom genomes can be constructed with the command line tool. GuideScan2 builds a memory-efficient compressed index based on Burrows-Wheeler Transform to annotate all sufficiently specific gRNAs, enabling multiple design and analysis tasks.

To illustrate the value of GuideScan2, we set out to assess the performance of gRNA libraries used in published CRISPR gene essentiality screens (**Fig. 2a-d, S1, S2**). Thanks to its novel design, GuideScan2 allows straightforward enumeration of off-targets and evaluation of the specificity of all gRNAs used in each screen, including gRNAs that are not part of the GuideScan2 database. Consistent with previous observations by us and others [3, 2, 4, 5, 6, 7], we found that a substantial number of gRNAs in published CRISPRko screens [8, 4, 9, 10, 11] have many off-targets and consequently low specificity (**Fig. 2d, S1a-d, Table S1**). GuideRNAs with particularly low specificity can confound CRISPRko essentiality screens by producing strong negative cell fitness effects even for non-essential genes (**Fig. 2a, S1e, Table S2**), likely through toxicity due to a large number of non-specific cuts. We found a similar bias in some CRISPR inhibition (CRISPRi) gene essentiality screens, where occasional low-specificity gRNAs were associated with false positive hits (**Fig. S2a**).

**Figure 2:**
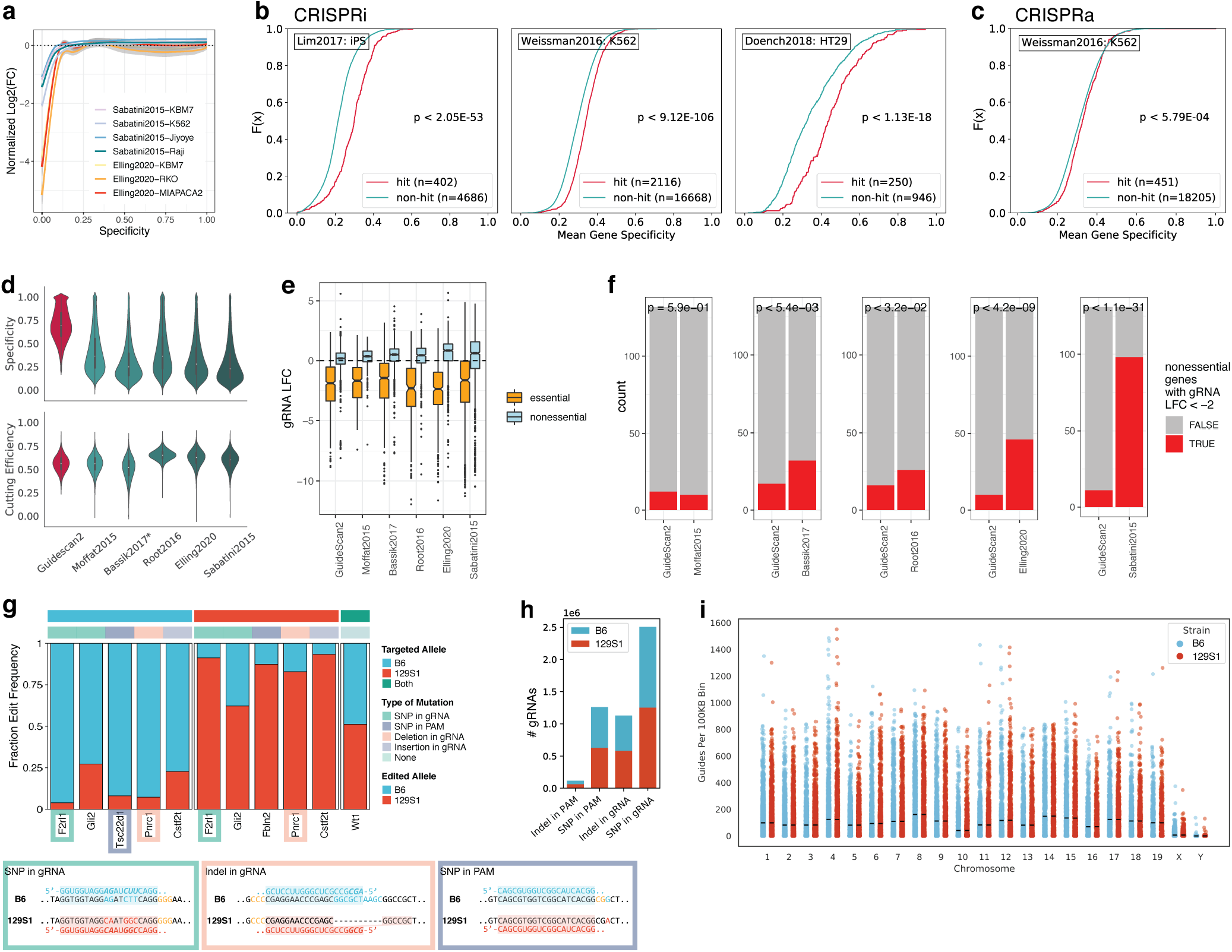
GuideScan2 reveals widespread confounding effects of low-specificity gRNAs in published CRISPR screens and enables design of new ready-to-use high-specificity gRNA libraries. (a) Reanalysis of published essentiality CRISPRko screens for non-essential genes. (b-c) Reanalysis of published (b) CRISPRi and (c) CRISPRa screens. Comparison of gene-level specificity (average over gRNAs per gene) for genes identified by the authors as hits and non-hits. (d) Comparison of new GuideScan2 human gRNA library against other libraries by specificity and cutting efficiency (average over gRNAs per gene). (e-f) Experimental results for the essentiality screen comparing new GuideScan2 gRNA library with other libraries. (e) gRNA targeting effect for essential and non-essential genes. (f) Fraction of non-essential genes in each pairwise library comparison that have at least one gRNA with strongly negative target effect (LFC *< −*2). (g) Experimental validation of selected C57BL/6- and 129S1/SvlmJ-specific gRNAs in mES cells of the F1 C57BL/6 *×* 129S1/SvlmJ hybrid mice. (h) Number of allele-specific gRNAs by category. (B6 stands for C57BL/6, 129S1 for 129S1/SvlmJ.) (i) Genome-wide frequency of C57BL/6- and 129S1/SvlmJ-specific gRNAs (excluding outliers, defined as bins with *>* 1700 gRNAs, total of 3 bins).

Importantly, we also found a different previously unobserved confounding effect of low gRNA specificity linked to *false negatives* in genome-wide CRISPRi and CRISPR activation (CRISPRa) screens. In such screens, multiple gRNAs per gene are used in order to increase statistical power and make up for occasionally problematic gRNAs. However, we observed that in published screens [12, 13, 14], genes identified by authors as the main hits tend to have significantly higher average gRNA specificity (**Fig. 2b,c, S2b-d**). This suggests that genes targeted by gRNAs with lower average specificity are systematically less likely to be called as hits of the screens, possibly because of dilution of targeting caused by excessive off-targets. In some cases, the predictive power of average gRNA specificity for calling genes as hits of the screen is comparable with that of the strongest biological factors identified by the authors (**Fig. S2d,e**). This newly identified confounding effect could present a major barrier for interpretation of results of genome-wide CRISPRi and CRISPRa screens.

To overcome potential problems with gRNA off-targets and low specificity in genome-wide screens, we designed new ready-to-use genome-wide CRISPR gRNA libraries targeting protein-coding genes in the mouse and human genomes, including six gRNAs per gene and complemented with safe-harbor-targeting and non-targeting gRNAs as controls (**Methods, Table S3**). gRNAs from our libraries have higher predicted specificity than other libraries while having similar cutting efficiency (**Fig. 2d, S1a, Table S1**). We expect that adoption of our GuideScan2 gRNA library for genome-wide CRISPR screens will help avoid the confounding effects of low-specificity gRNAs described above. Importantly, the strategy we used for designing this library (**Methods**) can be extended to design high-specificity gRNA libraries in various contexts across genomes, including for non-coding regions [2].

For experimental validation and comparison of the GuideScan2 library with five other gRNA libraries [8, 4, 9, 10, 11], we performed an essentiality screen in human A549 cells [15] (**Methods, Table S4**). The log fold change (LFC) in sequenced reads for each gRNA in this experiment measures the cell fitness effect from CRISPR editing with this gRNA, with large negative values indicating a more deleterious effect as expected for essential genes. As a positive control, we selected a random set of 100 essential genes and included all gRNAs for these genes from the six libraries into our screen. As expected, we observed that most of these gRNAs confer a strongly negative cell fitness effect that was comparable across libraries, confirming GuideScan2 efficiency (**Fig. 2e**). We also included negative control (non-targeting and safe-harbor-targeting) gRNAs in the experiment and confirmed that they do not result in lower cell fitness (**Fig. S3a, Methods**). The main arm of the experiment was designed to assess the effect of the lowest gRNA specificity on targeting non-essential genes. For each of the five libraries that we compared against GuideScan2, we selected the 140 potentially most problematic genes, defined as non-essential genes with the lowest gRNA specificity for that library, ranked by the average of the two lowest gRNA specificity values per gene. For each library our screen included all gRNAs for the lowest-specificity genes selected from this library, as well as all GuideScan2 gRNAs for the same genes, enabling direct pairwise comparison. As expected for non-essential genes, these gRNAs overall tended to have little effect on cell fitness, consistently across libraries (**Fig. 2e**). However, we observed that a fraction of low-specificity gRNAs in each library conferred a strong deleterious effect on cell growth (**Fig. S3b,c**). For four of the five libraries, the number of genes with lowest-specificity gRNAs was significantly higher than for GuideScan2, while for one library, Moffat2015 [8], the results were comparable with GuideScan2 (**Fig. 2f**). This is consistent with off-target analysis of the libraries performed using GuideScan2 where we found the Moffat2015 library to have the lowest number of off-targets (**Fig. S1b-d**). Overall, this suggests that GuideScan2 is effective at enumerating off-targets and estimating gRNA specificity, and this information can be used to reduce the confounding effects of low-specificity gRNAs by designing libraries that avoid such gRNAs.

To demonstrate GuideScan2’s flexibility in preprocessing and analyzing new genomes, we designed a genome-wide gRNA database for the F1 hybrid C57BL/6 *×* 129S1/SvlmJ mouse genome to enable allele-specific CRISPR targeting (**Fig. 2g-i, Table S5**). The database includes gRNAs that are expected to target one of the two alleles more efficiently, due to SNPs or indels in the target protospacer sequence or PAM or due to larger structural rearrangements in one of the chromosomes, while controlling for all potential off-targets in all chromosomes of the hybrid diploid genome (**Fig. 2g,h**). This database includes an approximately equal number of C57BL/6-specific and 129S1/SvlmJ-specific gRNAs, with an average of 92.6 C57BL/6-specific and 92.7 129S1/SvlmJ-specific gRNAs per 100Kb (**Fig. 2i**). For 33% of protein-coding genes, we found both at least one C57BL/6-specific and at least one 129S1/SvlmJ-specific exon-targeting gRNA. When selecting a subset of allele-specific gRNAs using the more stringent requirement of SNPs or indels in the PAM rather than in the protospacer sequence, we still observed broad genome-wide coverage by such gRNAs (**Fig. 2h**). We experimentally validated a number of allele-specific gRNAs and observed their high efficiency in allele-specific targeting (**Fig. 2g, Table S6**). In sum, this library is a convenient resource for allele-specific genome editing in F1 hybrid C57BL/6 *×* 129S1/SvlmJ mice. This example also demonstrates the power of GuideScan2 for gRNA design in custom settings.

In conclusion, GuideScan2 is a new flexible and efficient CRISPR gRNA design and analysis tool with command line and web interfaces that will facilitate CRISPR experiments across a wide range of applications.

**Figure S1:**
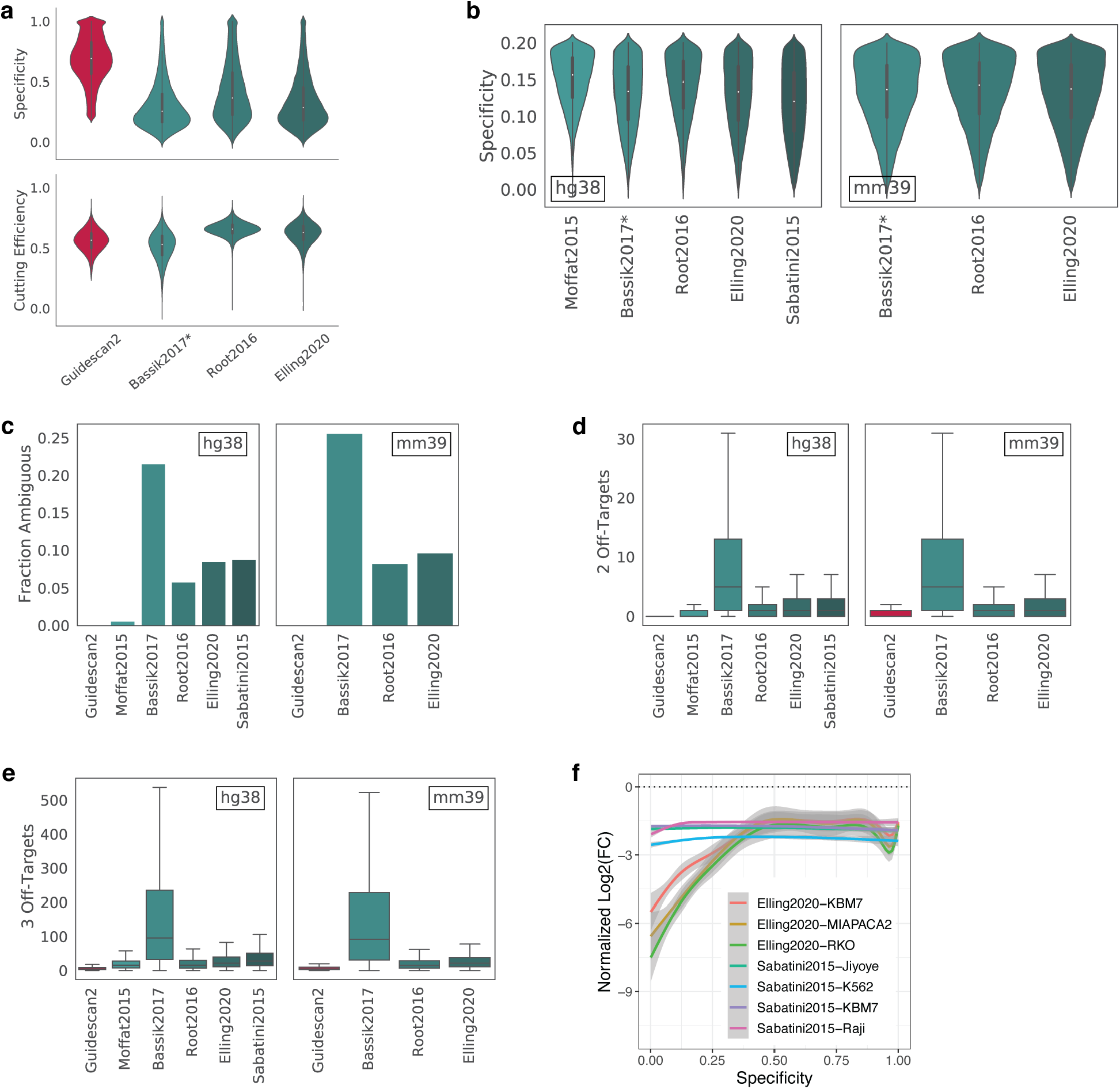
Reanalysis of published CRISPRko libraries. (a) Comparison of new GuideScan2 mouse gRNA library against other libraries by specificity and cutting efficiency (average over gRNAs per gene). (b) Distribution of specificity values (average over gRNAs per gene) at the range between 0 and 0.2 for human and mouse gRNA libraries. (c) Fraction of ambiguous genes across libraries. A gene is called “ambiguous” if there is a gRNA intended for that gene that has multiple perfect occurrences (without mismatches) in the genome. By design the GuideScan2 library does not have such gRNAs. (d) Boxplots (excluding outliers) of the number of 2-mismatch off-targets per gRNA. (e) Boxplots (excluding outliers) of the number of 3-mismatch off-targets per gRNA. (f) Reanalysis of published essentiality CRISPRko screens for essential genes.

**Figure S2:**
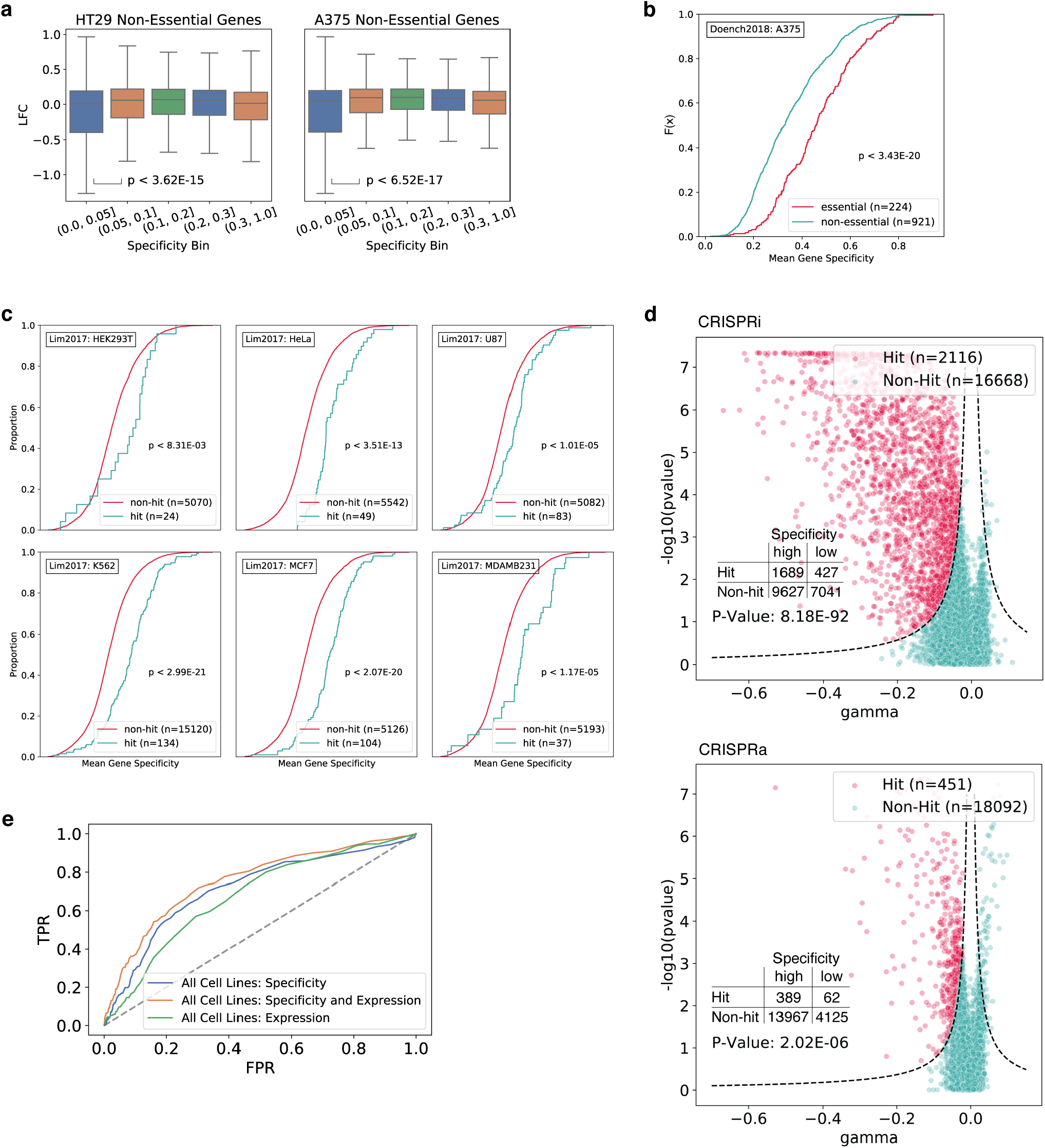
Reanalysis of published CRISPRi and CRISPRa libraries. (a) Reanalysis of the Doench2018 CRISPRi essentiality screen data. For non-essential genes, the gRNAs with specificity between 0 and 0.05 confer significantly lower LFC values than gRNAs with specificity between 0.05 and 0.1 (Mann–Whitney U test). (b) Comparison of gene-level specificity (average over gRNAs per gene) for genes identified by the authors as hits and non-hits in the Doench2018 CRISPRi essentiality screen. (c) Comparison of gene-level specificity (average over gRNAs per gene) for genes identified by the authors as hits and non-hits in the Lim2017 CRISPRi lncRNA screen across multiple cell lines. (d) Volcano plots for Weissman2016 CRISPRi/a screen showing the definitions of hits and non-hits from the original publication. *p*-value, Fisher’s exact test for enrichment of genes targeted with high-specificity (mean specificity *>* 0.25) gRNAs among hits. (e) ROC curve for logistic regression classifying between hits and non-hits using gRNA features in the Lim2017 lncRNA CRISPRi screen across all cell lines.

**Figure S3:**
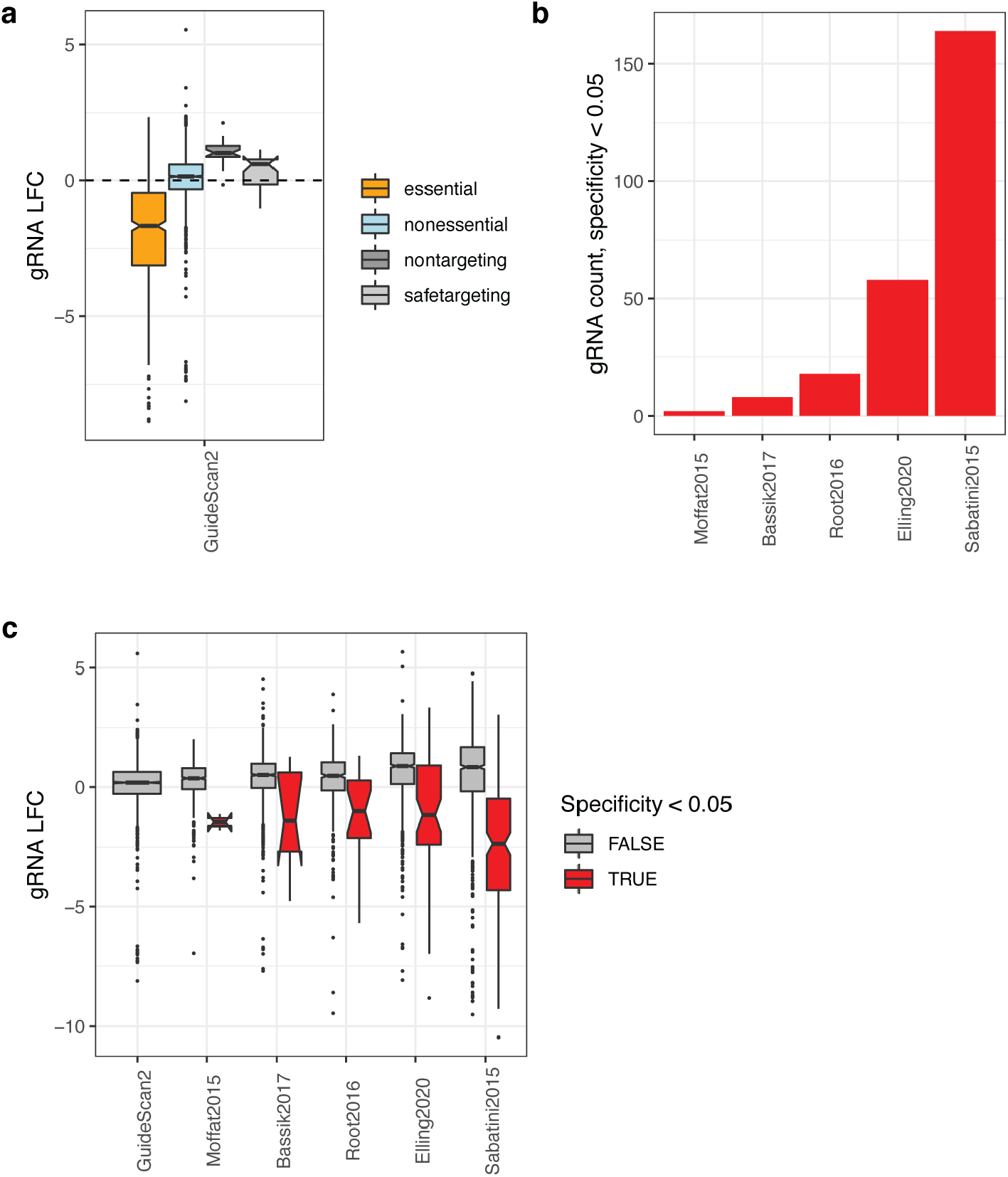
Essentiality screen for validation of GuideScan2 library. (a) GuideScan2 gRNA targeting effect for essential and non-essential genes and for non-targeting and safe-harbor-targeting controls. (b) Number of gRNAs with specificity below 0.05 in each library. By design GuideScan2 does not have gRNAs with specificity below 0.05. (c) Targeting effect for gRNAs depending on their specificity.

## Methods

### GuideScan2 algorithm

CRISPR-Cas systems can be used for targeted genome editing with RNA-guided endonucleases. They can be programmed to target a specific genomic location by designing a guide RNA (gRNA) with a particular short RNA sequence at its end, called the *spacer*. To elicit editing, the spacer sequence should be complementary or nearly complementary to the DNA sequence at the target location, called the *proto-spacer*, provided this sequence is adjacent to a particular protospacer adjacent motif (PAM). For example, CRISPR-Cas9 targets are generally thought to be determined by a 20-nucleotide spacer sequence at the end of the gRNA that is complementary to a DNA protospacer sequence followed immediately at the 3’ end by a PAM of the form NGG (resulting in more efficient targeting) or NAG (less efficient); here N stands for a ‘wildcard’, i.e. can match any nucleotide. Other natural and engineered CRISPR-Cas systems can vary in PAM sequence, PAM position with respect to the protospacer sequence, and requirements on the level of similarity between gRNA and the target. In what follows, unless otherwise noted, we will use CRISPR-Cas9 as our main example, but our approach is easily generalizable to other CRISPR systems. For simplicity we will associate the gRNA with its variable spacer sequence and will refer to it simply as the gRNA.

The task of gRNA design is, given a genomic region, to find gRNAs that can target anywhere in that region. Many potential gRNAs can target at multiple locations in the genome, though perhaps with varying efficiency. Typically a gRNA is designed to target a particular location with perfect complementarity. All other targets of this gRNA are then called *off-targets*. The goal of gRNA design is typically to maximize gRNA efficiency at the primary target site while minimizing off-targeting.

Variants and extensions of the gRNA design task include: paired gRNA design to select two gRNAs targeting flanking sites of a genomic region of interest; saturation experiment design to exhaustively select all gRNAs expected to target a selected region of interest; and library design to select a small number of the most effective gRNAs for each of hundreds or thousands of regions of interest.

To accomplish such tasks, our main approach is to preprocess the genome of interest to generate a genomic index. This genomic index can then be used for efficient queries reporting gRNA primary target and off-target information and for generation of the gRNA database for the given genome. The database contains gRNAs for the entire genome and information about off-targets of these gRNAs, according to a certain set of prespecified parameters, and provides the ability to efficiently search for gRNA information by genomic coordinates. The index can also be used to obtain additional information about gRNAs, e.g. about their more distant or non-conventional off-targets. It can also be used to search for information about any potential gRNAs, including those that were filtered out and excluded from the GuideScan2 database.

GuideScan used a retrieval tree (trie) data structure to preprocess the targetable space in the genome, i.e. all 20-mers followed by primary and secondary PAMs [3]. This trie was then used to construct a database, stored as a binary alignment map (BAM) file. This approach provided guarantees about identifying all potential off-targets of a gRNA up to a certain number of mismatches. However, searching for non-conventional off-targets or for information about gRNAs that were not included in the database (e.g. those with multiple perfect occurrences (without mismatches) in the genome) was hard because such queries required manipulation of the trie, which was excessively large.

GuideScan2 uses a data structure based on the Burrows-Wheeler Transform (BWT). BWT compresses long sequences while allowing for efficient search of short subsequences [16]. The BWT index has been used for building software packages for short-read alignment such as Bowtie and BWA [17, 18, 19]. However, we previously showed that these packages cannot be used for efficiently enumerating gRNA off-targets [3], since their implementation is specific for their task and not easily adaptable to the genome-wide search required for our analysis. GuideScan2 relies on an efficient implementation [20] of Wavelet Trees that utilize BWT based on rank-queries over the compressed sequence [21].

Our algorithm can be viewed as an evaluation of a single gRNA against ℬ, the BWT of a genome. That is, for a single gRNA with a spacer sequence *g* and PAM set P, for all *g*′ in a Hamming distance ball of radius *k* centered at *g*, we find all their occurrences in ℬ. We then validate these occurrences against the PAM set P, pruning any occurrences that are not followed by a PAM in this set. This set of validated occurrences forms the set of targets for this gRNA. Typically, we filter out gRNAs that have multiple perfect occurrences since we cannot distinguish the intended target, but this is a configuration parameter that can be modified, along with parameters *k* and P. The gRNA with a single perfect occurrence, considered to be its primary target, is then included in the database, and all other targets that contain mismatches are considered off-targets.

The aforementioned Hamming ball search is implemented as a depth-first search using rank-queries on the forward and reverse complement strands of the BWT of the genome. Because of the exponential blowup in search space even for small values of *k*, there are several heuristics applied to speed up the algorithm’s performance. First, we search for the reverse complement of the gRNA on the reverse complement strand so that the PAM validation, which typically contains an N, is done at the end of the search. This leads to an approximately four-fold performance improvement versus the naive strategy that searches for the forward guide on the forward strand, as the branching on N is done at the end when the search space has already been significantly pruned. Second, as mentioned earlier, we typically short circuit and stop searching when we find gRNAs with multiple perfect occurrences.

As a minor technical aside, we note that GuideScan2 is slightly more conservative than the original GuideScan tool when enumerating off-targets during database construction. Namely, unlike GuideScan, GuideScan2 does not penalize alternative PAMs. Thus, the set of off-targets produced by GuideScan2 is a strict superset of that output by the original Guidescan when running both tools with identical parameters. As in GuideScan, we define the gRNA specificity score using the formula from a previous publication [22] based on the CFD score [9]: Specificity is a value in the range [0, 1] that is closer to 0 for gRNAs with larger number of more closely matching off-targets. Importantly, the CFD score, and thus specificity, can only be calculated for gRNAs of length 20nt [9].

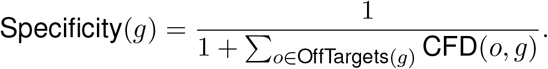

### Generation of genome-wide gRNA databases

We used GuideScan2 to generate a collection of gRNA databases (see http://guidescan.com) for frequently studied organisms for Cas9 and Cas12a (Cpf1) CRISPR systems. The databases are stored in a compressed BAM file format. Each row in the file corresponds to a gRNA and stores the genomic position and mismatch distance of all off-targets. Where applicable, the two scores *specificity* and *cutting efficiency* are stored in the `cs` and `ds` tags of the attributes field. Off-target information in these files is encoded in a hex format.

Database generation can be easily parallelized for efficiency. We took the following steps to parallelize the algorithm:

1. Generate the set of all 23-mers in the genome ending in NGG.
2. Partition this set of 23-mers into 1000 partitions.
3. Run GuideScan2 in parallel on each partition with alternative PAM NAG.
4. Merge the resultant SAM files together.
5. Append gRNA scores.
6. Convert SAM file to BAM format.

The database generation was performed using Memorial Sloan Kettering Cancer Center’s High Performance Computing resource. In step 5, we attached *cutting efficiency* score, defined using the Rule Set 2 [9], and *specificity*, as described above. This is done by a separate Python script that appends those scores to the SAM file. A complete tutorial describing the procedure for construction of genome-wide gRNA databases using GuideScan2 can be found in the software manual at https://github.com/schmidt73/guidescan-cli.

### CRISPRko screen analysis

We analyzed the relationship between gRNA specificity and performance in two published CRISPRko essentiality screens in human cell lines that we refer to as Sabatini2015 and Elling2020 [11, 10]. These are genome-wide essentiality screens where each gene is targeted with a small set of gRNAs and cell fitness depletion is measured for each gRNA. Specifically, each gRNA is associated with a control and treatment sequencing read count measured using RNA-seq, and from these counts fold-change values are computed. Genes with consistently strong depletion across the multiple gRNAs targeting them are thought to be essential in the modeled cellular context.

We preprocessed gRNAs from the Sabatini2015 and Elling2020 screens and converted the data into a unified CSV format. We used GuideScan2 (without off-target search) to verify the primary target locations of all gRNAs in these libraries. Human genome assembly hg38 was used as a reference. Then we ran GuideScan2 for all the gRNAs in these screens (including those not present in GuideScan2 database) against the hg38 GuideScan2 genomic index, searching for all off-targets with up to 3 mismatches while matching the alternative PAM sequence NAG.

Log2 fold change (LFC) values for the Elling2020 screens were taken from the publication [10]. LFC values for the Sabatini2015 screens were calculated from sequencing counts provided in the publication following the instructions in the publication [11]. Specifically, we filtered out any gRNAs with control counts below 400. We then normalized control and treatment counts by dividing by the total count in each column, respectively. These normalized counts were then used to compute LFC values. For the KBM7 cell line, the LFC values were averaged across the replicates.

To evaluate the relationship between gRNA specificity and its effect on cell fitness depletion in essentiality screens, we first classified gRNAs into two categories: those targeting essential and non-essential genes. For the Sabatini2015 data set, we chose a simple *p*-value threshold to classify genes into groups. Specifically, genes with all gRNAs with *p >* 0.01, as reported in the publication where *p*-values were calculated using a variant of the RNA interference gene enrichment algorithm (RIGER) [11], were classified as non-essential, while genes with all gRNAs with *p <* 0.01 were classified as essential. For the Elling2020 data, we used their published set of “Constitutive Core Essential Genes” and “Non Essential Genes” to classify gRNAs [10]. Regression between specificity and depletion using geom smooth() in R demonstrated that even for non-essential genes, gRNAs with particularly low specificity (*<* 0.15) have strongly negative LFC values comparable with those for essential genes (**Fig. 2a, S1f**).

We also set out to investigate how cell type-specific the gRNA effects are. To do this, we used the Sabatini2015 data since they targeted the same set of genes across four different cell lines. For each pair of cell lines, we computed the correlation between the LFC values across the same set of gRNAs and observed it to be less than 0.10 for every pair of cell lines. This suggests that gRNA cell fitness depletion effect is highly cell type-specific. Therefore the significant relationship between our GuideScan2-derived specificity score, calculated in a cell type-agnostic manner, and cell fitness depletion is especially surprising.

### CRISPRi and CRISPRa screen analysis

In order to investigate the effect of gRNA specificity in CRISPR inhibition and activation screens, we analyzed data from three publications herein referred to as Lim2017, Weissman2016 and and Doench2018 [12, 13, 14]. We preprocessed and reanalyzed gRNAs from these screens using GuideScan2 in the same manner as for CRISPRko screens. We thus obtained off-target information and specificity scores for all gRNAs in these screens.

The Lim2017 screen aimed to detect functional long non-coding RNAs (lincRNAs). In this screen, a set of 16,401 putative lincRNA transcription start sites (TSS) were targeted with ten gRNAs per site, and LFC depletion in the screen estimated for each gRNA. We observed across cell lines that lincRNAs reported to be functional, defined as hits of the screen as reported in the publication [12], had significantly higher mean gRNA specificity across the gRNAs targeting a lincRNA. This suggests that gRNA specificity can confound the interpretation of the results of these screens. Furthermore, extending the analysis in the publication [12], we performed binary logistic regression classification of genes to hits and non-hits, using data combined from all the seven cell lines. We used 80% of the data for training and 20% of the data for testing and reporting the performance. In the publication, the most predictive feature, by far, was gene expression (which is cell type-specific) [12]. Therefore in our analysis we compared specificity directly to gene expression. We discovered that the mean gRNA specificity outperformed gene expression (**Fig. S2e**) and that combining both features further improved predictive performance.

We performed similar analysis for the protein coding gene essentiality CRISPRi and CRISPRa screens Weissman2016 and Doench2018 [13, 14]. For the Weissman2016 data, we reproduced the definition of screen hits and non-hits using instructions from the publication [13] (**Fig. S2d**) and observed using Fisher’s exact test that genes with mean gRNA specificity above 0.25 were significantly enriched among hits. For this analysis, we only focused on “negative” hits. For the Doench2018 screen, we obtained the definitions of hits and non-hits from the publication [14].

### Reanalysis of genome-wide gRNA libraries

We reanalyzed previously published human and mouse genome-wide gRNA libraries from five publications [8, 4, 9, 10, 11] using GuideScan2. We refer to them as Moffat2015, Bassik2017, Root2016, Elling2020, and Sabatini2015, respectively. We observed that many gRNAs in these libraries had no-mismatch off-targets (i.e. multiple perfect occurrences in the genome) and overall low specificity (**Fig. 2d, S1**). Therefore we decided to construct new genome-wide gRNA libraries with high specificity, preserving and building on the design criteria of the previous libraries.

### GuideScan2 library design

We designed new genome-wide libraries using GuideScan2 and motivated by the design criteria of previous genome-wide libraries [8, 4, 9, 10, 11]. We constructed libraries for three reference genomes, hg38, mm10, and mm39, but analogous steps could be taken for any organism of interest.

The construction followed multiple steps as follows; the numbers of gRNAs after each filtering step are provided for the hg38 genome as an example. First, we identified the potential gRNA space as the set of all 20-mer sequences followed by an NGG PAM in the genome. Then, we filtered out gRNAs that have multiple occurrences in the genome with no mismatches or one mismatch, leaving us with 160,448,682 unique gRNAs. Then we further filtered out the gRNAs that do not cut within a CDS region, leaving us with 4,106,943 gRNAs. We further removed gRNAs with extreme G/C content, defined as having the total of C and G nucleotides greater than 80% or less than 20%, leaving us with 3,993,095 gRNAs. We then excluded any gRNAs containing monopolymers of length greater than three (i.e. four or more consecutive identical nucleotides), leaving us with 3,692,102 gRNAs in hg38. Finally, we filtered out gRNAs with cutting efficiency less than 0.25 or specificity less than 0.20. These remaining gRNAs were then used for our final library selection steps.

Using this filtered set of gRNAs, we attempted to find six gRNAs for each gene. For genes with six or fewer gRNAs, we used all of them. For genes with more than six gRNAs in the filtered set, we used further criteria to rank gRNAs and select six per gene. For this, we ranked gRNAs for each gene using a simple score that balances maximizing the gRNA specificity and cutting efficiency. Furthermore, often the nucleotide at the 5’ end of gRNA is replaced with a G for better efficiency, and therefore for each gRNA we consider specificity both for the gRNA itself and for a version of it where the nucleotide at 5’ end is replaced with a G (GuideScan2 allows to calculate both specificity values). To summarize, the score for ranking gRNAs for each gene is defined as

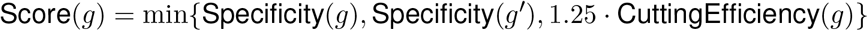

where *g*′ denotes the sgRNA *g* with the nucleotide at 5’ end replaced with a G. Cutting efficiency is defined as the “Rule Set 2” score [9]. For each gene that had more than six gRNAs in the filtered list, we selected six gRNAs with the highest value of Score().

Additionally, we included a set of 5,000 safe-targeting control gRNAs and 5,000 non-targeting control gRNAs. To generate non-targeting controls, we randomly generated gRNA sequences and selected those that had no genomic alignment with Hamming distance three or lower. For safe-targeting controls, we selected gRNAs that cut within the safe-targeting regions defined in “Supplementary Table 5” of the Bassik2017 paper [4]. As these safe-targeting regions were designed for hg19 and mm10 genome assemblies, we used LiftOver from the UCSC genome browser [23] to map the regions to hg38 and mm39, respectively. Among these safe-targeting guides, totalling 10,149,882 in hg38, we selected the 5,000 with the best specificity. For both sets, we applied the same filtering steps as for our gene-targeting gRNAs where applicable.

### Design of the essentiality screen for GuideScan2 library validation

In order to experimentally test the performance of our GuideScan2 genome-wide gRNA library and to confirm improved gRNA specificity without sacrificing efficiency, we designed an essentiality screen. The experiment consisted of two arms.

In the first arm, we screened a random subset of 100 essential genes as defined in a previous publication [24]. These 100 genes were selected from the top 5,000 most highly expressed genes (after averaging expression values over two replicates), defined using published RNA-seq gene expression data. For these genes, we included all gRNAs from the five libraries and from the GuideScan2 library targeting these genes, resulting in 4,050 gRNAs in this arm of the screen.

The second arm of the experiment aimed to demonstrate decreased confounding effect of low-specificity gRNAs in our GuideScan2 library. For this we used gRNAs targeting non-essential genes. Namely, we hypothesized that low-specificity gRNAs in other libraries reduce cell fitness even when targeting non-essential genes and thus confound the interpretation of results of screens relying on these libraries. To test this, we first selected a set of non-essential genes from Hart et al. [25] that were poorly expressed in the A549 cell line. For all libraries except Bassik2017, we ranked genes by the average specificity of their two lowest specificity gRNAs. Then we included all gRNAs targeting the 100 top ranking (lowest specificity) genes for each library in our screen. For Bassik2017 we applied a slightly different ranking scheme as the specificity score is not defined for gRNAs of length other than 20nt. We ranked genes by the average number of off-targets for the two gRNAs with the largest number of off-targets. For comparison to the GuideScan2 library, we included all GuideScan2 library gRNAs targeting the same genes. Then we selected the 100 genes with the highest such rank. This second arm of the screen totaled 7,130 gRNAs targeting 400 distinct genes.

Finally, we included a random set of 100 safe-targeting and 100 non-targeting control gRNAs from the GuideScan2 library into our screen.

This resulted in the total of 10,033 unique gRNAs selected for the screen.

### CRISPR-Cas9 essential genome screen

#### Cloning of the CRISPR libraries

sgRNAs were divided into small pools of libraries, and oligo pools were synthesized by Agilent Technologies. To screen these libraries, we generated the pUSEBR (U6-sgRNA-EFS-Blast-P2A-TurboRFP) lentiviral vector by Gibson assembly of the following DNA fragments: (i) PCR-amplified U6-sgRNA (improved scaffold) [26] cassette, (ii) PCR-amplified EF1a promoter, (iii) PCR-amplified Blast-P2A-TurboRFP gene fragment (IDT), and (iv) BsrGI/PmeI-digested pLL3-based lentiviral backbone [27]. Libraries were cloned into pUSEBR using a modified version of the protocol published by Doench et al. to ensure a library representation of *>* 10,000-fold. Briefly, each library was selectively amplified using uniquely barcoded forward and reverse primers that append cloning adapters at the 5’ and 3’ ends of the sgRNA insert, purified using the QIAquick PCR Purification Kit (Qiagen), and ligated into BsmBI/Esp3I-digested and dephosphorylated pUSEBR, using high-concentration T4 DNA ligase (NEB). A total of 1.2 µg of ligated pUSEBR-CRISPR Library plasmid DNA was then electroporated into Endura electrocompetent cells (Lucigen). Competent cells were recovered for 1 h at 37ºC, plated across four 15-cm LB-Carbenicillin plates (Teknova), and incubated at 37ºC for 16 h. The total number of bacterial colonies per sub-pool was quantified using serial dilution plates, to ensure a library representation of *>* 10,000-fold. The next morning, bacterial colonies were scraped and briefly expanded for 4 h at 37ºC in 500 mL of LB-Carbenicillin. Plasmid DNA was isolated using the QIAfilter Plasmid Maxi Kit (Qiagen). To assess sgRNA distribution, the sgRNA target region was amplified using primers that append Illumina sequencing adapters on the 5’ and 3’ ends of the amplicon, as well as a random nucleotide stagger and unique demultiplexing barcode on the 5’ end. Library amplicons were size-selected on a 2% agarose gel, purified using the QIAquick Gel Extraction Kit (Qiagen), and sequenced on an Illumina NextSeq instrument (75 nt single-end reads).

#### CRISPR-Cas9 screening

A549 cells expanded from a single clone harboring a doxycycline-inducible Cas9 (Addgene #114010) were a gift from J.T. Poirier [15]. To ensure that most cells harbor a single sgRNA integration event, the volume of viral supernatant that would achieve an MOI of 0.1 upon transduction was determined. Briefly, cells were plated at a density of 0.5×10^6^ cells per well in 12-well plates along with increasing volumes of a 1 to 10 dilution of the master pool viral supernatant and polybrene (EMD Millipore, 8 µg/mL). Cells were then incubated at 37°C overnight. Viral infection efficiency was determined by the number of surviving blasticidin-resistant (Gibco, 10 µg/mL) cells as compared to unselected control assessed using Countess II FL (Thermo Fisher). The volume of viral supernatant that achieve 10% infection rate was used in the screen. To ensure a representation of 3250X at the transduction step, the appropriate number of cells were transfected with viral supernatant in 150 mm tissue culture plates (Corning) per infection replicate and selected with blasticidin for 5 days. Subsequently, half of the blasticidin-selected cells were pelleted and stored at -80°C (cumulative population doubling T0/Input population) while the rest were plated and treated with doxycycline (Millipore-Sigma, 1 µg/mL) for 18 days, or approximately 18 population doublings (TF/Final). At least 200 million cells were harvested and pelleted for this final time point.

#### Genomic DNA isolation

Genomic DNA was extracted from cells using phenol-chloroform extraction. Briefly, cell pellets were incubated at 50°C overnight in lysis buffer (100mM Tris-HCl pH 8.5, 200mM NaCl, 5mM EDTA, 0.2% SDS) containing 200 µg/mL proteinase K (Roche). After digestion, the supernatant was first extracted using phenol:chloroform:isoamyl alcohol 25:24:1 pH 8.0 (Millipore-Sigma), followed by chloroform:isoamyl alcohol 49:1 (Millipore-Sigma). The gDNA was then precipitated with equal volume of 100% isopropanol, washed with 70% ethanol, and resuspended in UltraPure distilled water (Gibco).

### Library amplification of CRISPR screens

Libraries were amplified from gDNA by a modified 2-step PCR version of the protocol published by Doench et al. Briefly, an initial “enrichment” PCR was performed, whereby the integrated sgRNA cassettes were amplified from gDNA (PCR#1), followed by a second PCR to append Illumina sequencing adapters on the 5’ and 3’ ends of the amplicon, as well as a random nucleotide stagger and unique demultiplexing barcode on the 5’ end (PCR#2). Each “PCR#1” reaction contained 25 µL of Q5 High-Fidelity 2X Master Mix (NEB), 2.5 µL of Nuc PCR#1 Fwd Primer (10 µM), 2.5 µL of Nuc PCR#1 Rev Primer (10 µM), and 5 µg of gDNA in 20 µL of water. PCR#1 amplicons were selected on a 2% agarose gel and purified using the QIAquick Gel Extraction Kit (Qiagen). These amplicons were then used as template for “PCR#2” reactions. Each PCR#2 reaction contained 25 µL of Q5 High-Fidelity 2X Master Mix (NEB), 2.5 µL of a unique Nuc PCR#2 Fwd Primer (10 µM), 2.5 µL of Nuc PCR#2 Rev Primer (10 µM), and 300 ng of PCR#1 product in 20 µL of water. Library amplicons were size-selected on a 2% agarose gel, purified using the QIAquick Gel Extraction Kit (Qiagen), and sequenced on an Illumina NextSeq500 instrument (75 nt single end reads). PCR settings for PCR#1 and PCR#2 were: initial denaturation at 98°C for 30 s; then 98°C for 10 s, 65°C for 30 s, 72°C for 30 s for 24 cycles; followed by extension at 72°C for 2 min.

### Analysis of CRISPR-Cas9 screen data

FASTQ files were processed and aligned to the reference sgRNA library file, and log2 fold change value for each sgRNA was calculated using MAGeCK [28]. Only the gRNAs with control read count *>* 100 were used in subsequent analysis. For analysis of gRNAs targeting essential genes, average LFC per gene was calculated across all gRNAs used in the screen for that gene, and only the genes with such average below -1 were used, resulting in a selection of 85 genes out of 100 genes. The remaining analysis for both essential and non-essential genes followed in a straightforward way from the design of the screen.

### Allele-specific CRISPR-Cas9 editing

#### Design of allele-specific sgRNAs

We used GuideScan2 to design a set of allele-specific gRNAs for an F1 cross between C57BL/6 and 129S1/SvlmJ mouse strains, thereafter referred to as B6 and 129S1.

First, we constructed a synthetic 129S1 pseudo-genome using MMARGE [29] by introducing sequence variants obtained from the Mouse Genome Project (`files 129S1_SvImJ.mgp.v5.snps.dbSNP142.vcf` and `129S1_SvImJ.mgp.v5.indels.dbSNP142.normed.vcf` obtained from ftp://ftp-mouse.sanger.ac.uk/REL-1505-SNPs_Indels/strain_specific_vcfs) [30] to the reference B6 genome assembly mm38 (file `GRCm38_68.fa`). Then we constructed the GuideScan2 gRNA database for this synthetic 129S1 pseudo-genome, as well as separately for the B6 genome.

For each gRNA from the 129S1 database, we used GuideScan2 to search for it in the B6 index and filtered it out if a perfect occurrence was found; we also performed a reverse analysis and filtering of B6 gRNAs against the 129S1 index. The remaining gRNAs after this filtering formed the list of allele-specific gRNAs. We then used GuideScan2 to search for off-targets of these gRNAs in both 129S1 pseudo-genome and B6 genome.

The allele-specific gRNAs were annotated with protein-coding genes from GENCODE vM29 (file `gencode.vM29.annotation.gtf.gz`). A gRNA was annotated with a gene if the corresponding cutting site (3nt upstream of PAM) was within an exon of that gene. If multiple such gene annotations satisfied this criterion, an arbitrary one of them was chosen. Of the 21,812 protein-coding genes, 8,079 had a B6-specific gRNA and 8,330 had a 129S1-specific annotation, while 7,187 genes had both.

A representative set of sgRNAs were selected from the allele-specific database for the F1 B6/129S1 cross described above. crRNA, according to the differential sgRNA spacer sequences, were synthesized by IDT.

### CRISPR-Cas9 editing

V6.5 mouse embryonic stem cells (gift from Rudolf Jaenisch), which was generated from a F1 cross between C57BL/6 and 129S1/SvlmJ mouse strains, were plated in 6-well plates at a density of 0.5×10^6^ cells per well 24 hours prior to transfection. Alt-R CRISPR-Cas9 crRNAs with spacer sequences that would differentially target the C57BL/6 and 129S1/SvlmJ alleles, as well as a control spacer that would not differentially target the C57BL/6 and 129S1/SvlmJ alleles, were synthesized by IDT and were annealed with Alt-R CRISPR-Cas9 tracrRNA ATTO 550 (IDT) according to manufacturer’s protocol. Following, Alt-R S.p. Cas9 nuclease V3 (IDT) was mixed with the crRNA-tracrRNA duplex and transfected into V6.5 mESCs using Lipofectamine CRISPRMAX (Thermo Fisher) according to manufacturer’s protocol. Transfection efficiency was determined by ATTO 550 fluorescence after 24 hours, as visualized by the Axio Observer A1 inverted microscope (Zeiss). Cells were then collected at 72 hours post transfection and genomic DNA was extracted using phenol-chloroform as previously described above.

### CRISPR sequencing

Edited gDNA sequences were amplified using Q5 polymerase (NEB) using specific PCR primer pairs that flank an approximately 250bp region encompassing the CRISPR-Cas9 cut site. Each reaction contained 10 µL of Q5 Reaction Mix (NEB), 10 µL of Q5 GC Enhancer Buffer (NEB), 2.5 µL of a forward primer (10µM), 2.5 µL of a reverse primer (10 µM), 1µL of dNTPs (10mM), 0.5µL of Q5 polymerase (NEB), and 1µg of gDNA in 23.5µL of water. PCR settings were: initial denaturation at 98°C for 3 min; then 98°C for 10 s, 65°C for 30 s, 72°C for 30 s for 25 cycles; followed by extension at 72°C for 2 min. PCR amplicons were size selected on a 2% agarose gel and purified using the QIAquick Gel Extraction Kit (Qiagen), and sequenced and deconvoluted using the Amplicon-EZ service provided by GENEWIZ/AZENTA (South Plainfield, NJ).

### Analysis of CRISPR-Cas9 editing data

Sequence abundance files provided by GENEWIZ/AZENTA were analyzed for allele-specific editing using custom Python 3 scripts. Briefly, each unique sequencing read was identified as either C57BL/6, 129S1/SvlmJ, or ambiguous based on the presence of sequence variations caused by SNPs or indels that were outside of the cut site. Ambiguous reads were discarded, and cumulative fractions of reads attributed to C57BL/6 or 129S1/SvlmJ were then calculated and tabulated for each unique spacer sequence.

## Supplementary tables

**Table S1**: Parameters of GuideScan2 and other published genome-wide gene-targeting gRNA libraries.

**Table S2**: Reanalysis of gRNA specificity in published CRISPR screens.

**Table S3**: GuideScan2 genome-wide gene-targeting gRNA libraries for mouse and human.

**Table S4**: Design and results of the validation gene essentiality screen of the human GuideScan2 gene-targeting library.

**Table S5**: Genome-wide gRNA library for allele-specific targeting in F1 C57BL/6 *×* 129S1/SvlmJ hybrid mice.

**Table S6**: Design and results of the allele-specific targeting validation experiment.

## Acknowledgements

We thank Marco Russo, Ralph Garippa and the Gene Editing and Screening Core Facility at MSKCC, Christof Fellmann, Alex Perez, and Joana Vidigal for helpful suggestions about the manuscript and the GuideScan2 web interface, and J.T. Poirier for providing the A549 cells. This work was supported by Geoffrey Beene Cancer Foundation, MSK Functional Genomics Initiative, Agilent Technologies Thought Leader Award (SWL), NIH grants U01 HG009395 (CSL and AV), U01 HG012103 (CSL), 5R01CA23412 (AV), and P01 CA129243 (SWL). F.J.S.-R. was supported by the MSKCC TROT program (5T32CA160001), a GMTEC Postdoctoral Researcher Innovation Grant, and is an HHMI Hanna Gray Fellow. S.W.L. is the Geoffrey Beene Chair of Cancer Biology and an Investigator of the Howard Hughes Medical Institute.

## Disclosures

S.W.L. is an advisor for and has equity in the following biotechnology companies: ORIC Pharmaceuticals, Faeth Therapeutics, Blueprint Medicines, Geras Bio, Mirimus Inc. and PMV Pharmaceuticals. S.W.L. acknowledges receiving funding and research support from Agilent Technologies for the purposes of massively parallel oligo synthesis.

## Notes

https://guidescan.com/

